# Coevolution increases robustness to extinctions in mutualistic but not antagonistic communities

**DOI:** 10.1101/2023.05.22.541738

**Authors:** Fernando Pedraza, Klementyna A. Gawecka, Jordi Bascompte

## Abstract

Co-extinctions may exacerbate the current biodiversity crisis. Yet, we do not understand all the factors that shape the robustness of communities to the loss of species. Here we analyse how coevolution influences the robustness of mutualistic and antagonistic communities. We find that coevolution increases robustness in mutualism but reduces it in antagonism. These differences are due to coevolution altering the density of interactions in communities. The largest changes to robustness occur when coevolutionary selection is strong. Yet, the effect size of coevolution on robustness depends on the size of the community. Our results may broaden the suite of mechanisms affecting the resilience of ecological communities. These insights may inform efforts to reduce the risk of species loss in the face of global change.

## Introduction

Species interactions have the potential of amplifying extinction events (Dunn et al., 2009; Colwell et al., 2012; Brodie et al., 2014). Co-extinctions occur when the extinction of one species causes the loss of other dependent species. For instance, the extinction of a host triggers the co-extinction of its parasites (Stork and Lyal, 1993; Gompper and Williams, 1998). Yet, beyond pairwise associations, species constitute networks of interactions. The structure of such interaction networks, in turn, affects the likelihood of co-extinctions (Solé and Montoya, 2001; Memmott et al., 2004). For example, networks with a large density of interactions (i.e. connectance) tend to be more robust to secondary extinctions than poorly connected ones (Dunne et al., 2002; Eklö f and Ebenman, 2006, but see Vieira and Almeida-Neto (2015); Morrison et al. (2020)).

The structure of interaction networks varies in time and space (Guimarães, 2020). These changes are caused by the interplay between the environmental conditions and biological processes mediating interactions (Mittelbach and Schemske, 2015; Poisot et al., 2015; Guimarães, 2020). Turning to the biotic drivers, the behaviour, abundance, and traits of species can all play a role in determining interactions (Vázquez, 2005; Santamaŕıa and Rodŕıguez-Gironés, 2007; Vázquez et al., 2009; Kaiser-Bunbury et al., 2010; Gibert and DeLong, 2017). For instance, variation in traits and changes in foraging behaviour can alter the structure of food webs (Brose et al., 2019) and plant-pollination networks (Spiesman and Gratton, 2016). These biotic drivers are, themselves, the result of ecological and evolutionary processes. Thus, the assembly of interaction networks is shaped by the ecology and evolution of interacting species (Post and Palkovacs, 2009; Mittelbach and Schemske, 2015; McPeek, 2017; Ponisio et al., 2019).

Coevolution leads to the reciprocal evolutionary change of interacting species. Often, these coevolutionary changes occur in the traits that mediate interactions (Thompson, 2005). For example, coevolution explains the covariation in fly proboscis length and the tube length of the flowers they pollinate (Anderson and Johnson, 2008). As a result, coevolution potentially influences the establishment and maintenance of interactions (Thompson, 2005; Santamaŕıa and Rodŕıguez-Gironés, 2007; Vázquez et al., 2009; Guimarães, 2020). In fact, theoretical work suggests that coevolution can give rise to networks that resemble empirical ones (Valverde et al., 2018; Andreazzi et al., 2020). Furthermore, the intensity of coevolution can influence network structure. In this sense, Nuismer et al. (2013) showed that the strength of coevolutionary selection modulates the density of interactions of mutualistic networks. Yet, we ignore how coevolutionary selection influences network structure beyond mutualistic interactions. Moreover, we lack a general understanding of how coevolution, by affecting network structure, influences the spread of co-extinctions.

Here we explore the role of coevolution in shaping the robustness of mutualistic and antagonistic communities to secondary extinctions (Figure 1). We first simulate the assembly of interaction networks based on the coevolution of species traits. We then characterise their structure and quantify their robustness to secondary extinctions. Finally, we compute the difference in network robustness between randomly assembled networks and those built by coevolution. Throughout, we highlight differences between i) mutualistic and antagonistic communities and ii) different intensities of coevolution.

**Figure 1:**
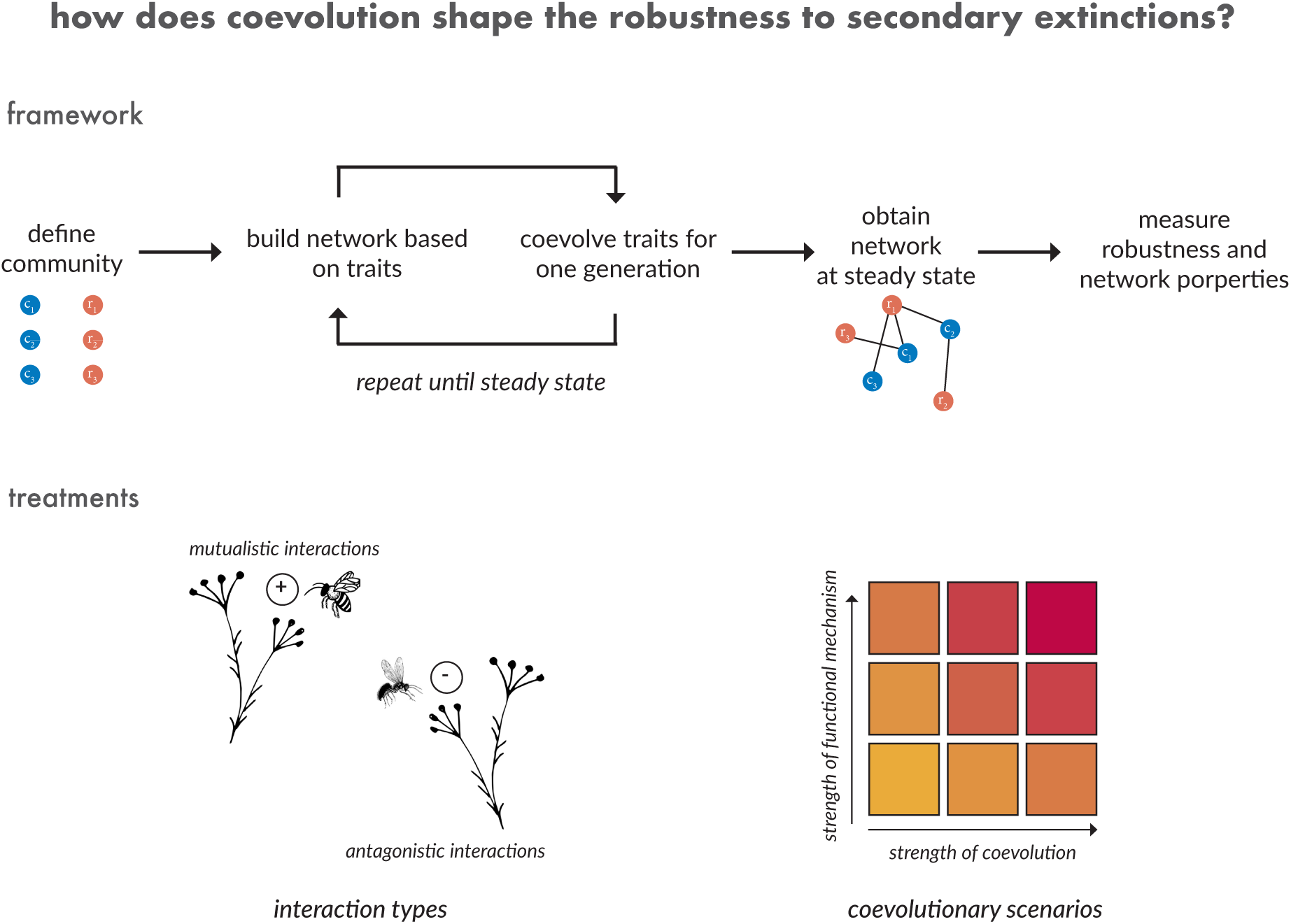
Outline of framework and experimental treatments. To test how coevolution shapes the robustness to secondary extinctions, we first define a community by: (i) specifying the number of consumer (c) and resource (r) species in each guild and (ii) assigning each species an initial trait value and environmental optimum. Then, we (iii) build a network of interactions based on species’ similarity of traits and (iv) let species traits coevolve for one generation. We repeat steps (iii) and (iv) until we reach a steady state —when trait values and network structure no longer change between timesteps. Lastly, we (v) measure the robustness to secondary extinctions and the structure of the interaction networks obtained. We perform simulations for: communities with antagonistic or mutualistic interactions and contrasting coevolutionary scenarios.

## Methods

### Modelling framework

Our modelling framework (Figure 1) allows us to simulate the coevolution of mutualistic or antagonistic communities. We next describe the three components of the modelling framework.

### Defining communities

Each community is defined be three properties. i) The number of species in each guild. We assume that communities are composed of two guilds: plants and animals. We hereafter refer to plants as resources and animals as consumers regardless of the type of interaction they engage in. ii) The type of interactions in the community. We assume that communities are composed entirely of either mutualistic or antagonistic interactions. Interaction types are fixed and do not change over time. iii) The environmental optimum (*θ*) and initial trait value (*Z^t^*^0^) of each species in the community. We assume that each species possess a single quantitative trait (*Z^t^*) and a corresponding optimal value of this trait (*θ*). For each species, we draw a random number from a given distribution (see simulations section) and use it to define both the environmental optimum value and initial trait value of each species. We assume that species’ traits at the start of the simulation are equal to their environmental optimum. We draw *θ* and *Z^t^*^0^ from different distributions as part of our effort to test how the niche breadth of a community affects the building of species interaction networks (see simulations section).

### Building networks

We build networks based on species traits. We assume that species with more similar traits are more likely to interact. We define the probability of a pair of species (*i*, *j*) interacting as:

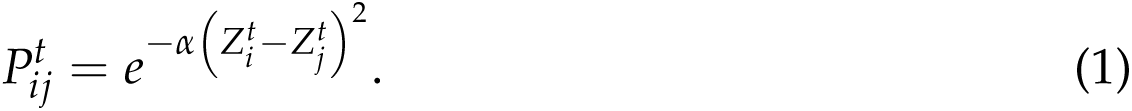

Where 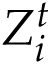 and 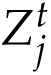 are the traits of species *i* and *j* at time *t*. *α* (hereafter strength of functional mechanism) is a constant that determines the sensitivity to trait matching. As *α* increases, species’ traits have to be increasingly similar in order to maintain a high interaction probability. We vary *α* as part of our effort to test how changes in the way coevolution operates affect the evolution of species interaction networks.

We use Equation 1 to calculate the interaction probability of all pairwise combinations of consumers and resources. Then, for each species, we find the pairwise combination with the highest probability and establish it as an interaction. By doing so, we guarantee that all species in the community have at least one interaction. Next, for all remaining pairwise combinations, we draw a random number *r* on the interval [0, 1). We assume that an interaction between a pair of species only occurs when 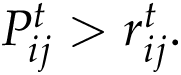.

We do not set an upper bound to the number of interactions that can be established. Thus, the connectance of each network (i.e. the proportion of realised interactions from the pool of all possible interactions between species) emerges as a result of the coevolution of traits.

### Simulating coevolution

We model species coevolution using the frameworks proposed by Guimarães et al. (2017) and Andreazzi et al. (2017, 2020), for mutualism and antagonism, respectively. These models use a selection gradient approach to track the change of a single quantitative trait for each species in a community across discrete time as a result of the selection imposed by interactions and the environment. We define the mean trait evolution of species *i* over a time step as:

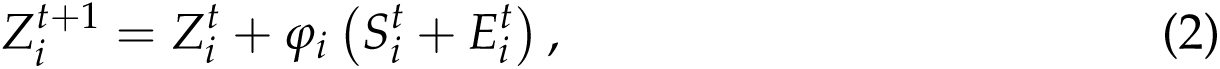

where *φ_i_* is a compound parameter that affects the slope of the selection gradient and is proportional to the additive genetic variance, while 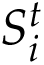 and 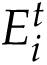 are the partial selection differentials attributed to selection imposed by interactions and environment, respectively.

Starting with how interactions shape species evolution, we assume that the extent to which species traits change as a result of interactions depends on the degree of trait similarity between interacting species:

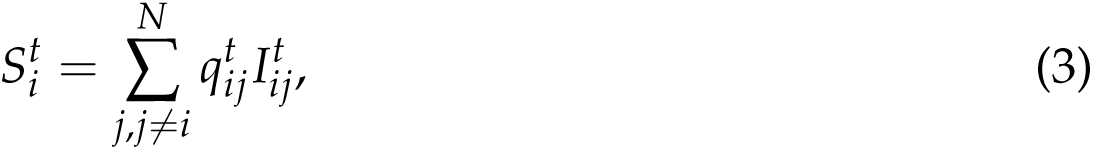

where *N* is the number of species in the network and 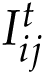 is the trait value selected by the interaction of species *i* with species *j*, and depends on the type of interaction (either mutualistic or antagonistic, see details below). 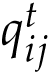 describes the evolutionary effect of species *j* on species *i* and serves to weigh the relative importance of the selection imposed by species *j* to the selection gradient compared with all other sources of selection. We define 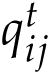

as:

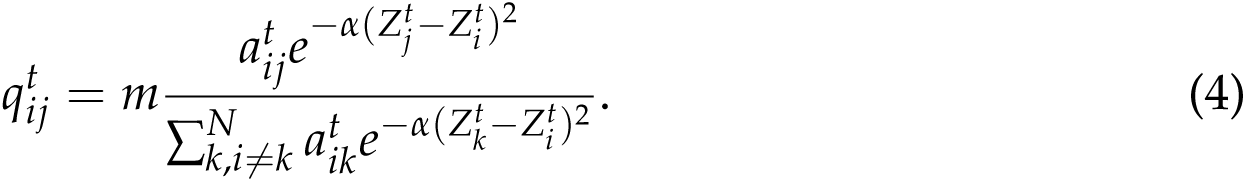

*m* is the strength of coevolutionary selection (hereafter ‘strength of coevolution’). It captures the relative importance of interactions in shaping trait evolution. At the extremes, trait evolution is completely driven either by the environment (*m* = 0) or by the interactions (*m* = 1). 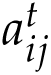 is an element of the symmetric binary adjacency matrix, *A^t^*, of the network (where 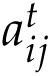 = 1 if species *i* and *j* interact, and 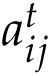 = 0 otherwise). *α*, as before, is a constant that determines the sensitivity to trait matching.

We assume that the phenotype selected by the interaction of species *j* with *i* (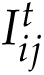in Eq. 3) depends on the type of interaction.

For mutualistic interactions, we assume that selection imposed by partners favours trait matching. Phenotypic matching in mutualism is well described and may be necessary for the successful interaction of mutualistic partners (Jordano et al., 2003; Agosta and Janzen, 2005). For example, in the matching between the depth of floral corolla and the length of hummingbird bills (Dalsgaard et al., 2008) or the size of seeds and body mass of frugivores (Jordano, 1995).

We assume that species with more similar traits interact more frequently and thus, exert strong reciprocal selection on each other. Consequently, species with more dissimilar traits interact less often and exert weaker reciprocal selection on each other. We define the effect of species *j* on species *i* at time *t* as:

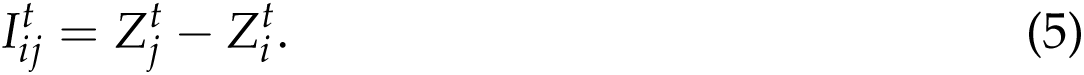

For antagonistic interactions, we assume that selection imposed by interactions favours trait similarity for consumers and mismatch for resource species. This assumption captures systems where the chance of a successful attack of a consumer on its resources increases when its traits match the defences of its victims, while the chances of a resource escaping its consumer increases if its traits mismatch those of its consumer (Nuismer and Thompson, 2006; Hanifin et al., 2008). Such dynamics are observed in the interaction between hard-shelled invertebrates and their shell-destroying predators (Vermeij, 1994) or between crustaceans and their parasitic bacteria (Luijckx et al., 2013).

For consumers, the effect of resource *j* on consumer *i* at time *t* is defined by Eq. 5. For resources, trait mismatch results in its traits either increasing (if 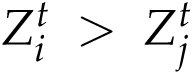) or decreasing (if 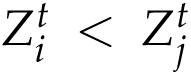). A critical mismatch value *ε* determines whether traits increase or decrease (Andreazzi et al., 2017, 2020)). If 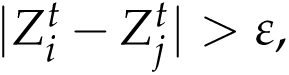 then the consumer has a negligible effect on the resource 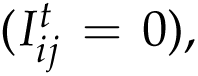, as in commensalism. Otherwise, if 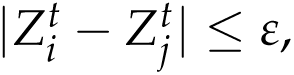 the effect of consumer *j* on resource *i* at time *t* is expressed as:

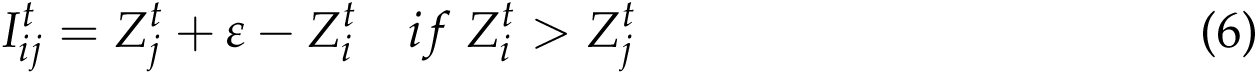

or

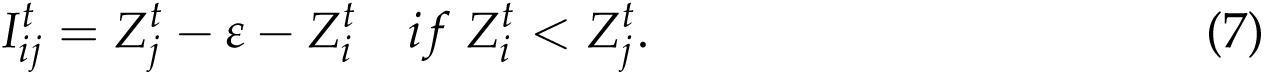

Finally, we define the trait change imposed by the environment, 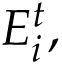 as:

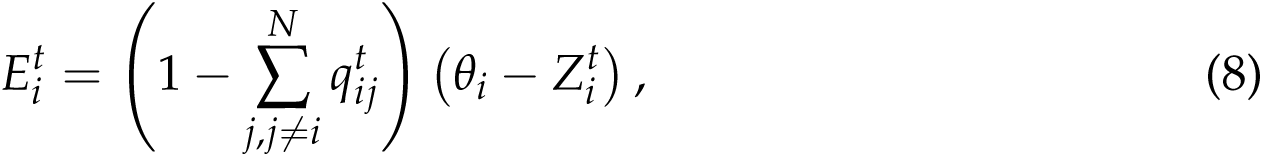

where *θ_i_*is the environmental optimum of species *i* (i.e., the phenotype favoured by the environmental selection).

### Simulation setup

To test how coevolution shapes the robustness of communities to secondary extinctions, we use the coevolutionary framework to construct interaction networks. To do so, we begin by defining communities (see corresponding section). Next, we build an initial network of interactions based on the trait values of the species that make up the community (Equation 1) and let these traits coevolve for one timestep (Equation 2). We then repeat the process of building the interaction network and allowing traits to coevolve at each timestep for a total of 200 timesteps. This provides sufficient time for the network of interaction and the species traits to reach a steady state, where trait values and network structure no longer change between timesteps. At timestep 200, we store the network of interactions for further analysis (Figure 1).

We performed the above simulations for: (i) communities with either mutualistic or antagonistic interactions and (ii) communities that experience different coevolutionary scenarios. A coevolutionary scenario is defined by a particular combination of the strength of coevolution (*m*) and the strength of functional mechanism (*α*). We vary the strength of coevolution from 0.05 to 0.95 and the strength of functional mechanism from 0.01 to 1. In total we test 399 coevolutionary scenarios (19 *m* values x 21 *α* values).

We constructed interaction networks for 30 communities. These communities differed in their size and ratio of consumers to resources (Figure S1). However, to facilitate the comparison between the mutualistic and antagonistic coevolution treatments, we used the same communities for both interaction types. Yet, we also constructed interaction networks for 30 empirical mutualistic and antagonistic communities archived in the Web of Life repository (http://www.web-of-life.es/, Fortuna et al., 2014). We leverage these to test whether our modelling framework was able to recreate the observed connectance values of the empirical networks. We found that for the parameter space we explored, our approach was able to recreate the observed connectance of antagonistic communities better than mutualistic ones (Figure S2 left panel). Furthermore, the coevolutionary scenarios that best predicted the observed connectance differed between interaction types. Mutualistic networks were best reproduced with high *α* values, whereas antagonistic ones with low *α* and *m* values (Figure S2 right panel).

To test the sensitivity of our findings, we performed simulations for communities with broad or narrow niches. For broad niche communities, the initial trait values 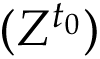 and environmental optima of species (*θ*) were drawn from a uniform distribution (*U*_[0,10]_). For narrow niche communities, the initial trait values 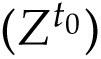 and environmental optima of species (*θ*) were drawn from a truncated normal distribution (*N*_(5,1)_ on the interval [0, 10]). Since broad niche communities performed best in our empirical validations (Figure S2 left panel), we present results from these communities in the main text and show results for narrow niche communities as supplementary material (Figures S3-S10).

Lastly, for each community, we performed 30 replicate simulations for each combination of interaction type, coevolutionary scenario, and niche breadth. The replicate simulations differed in the sampled 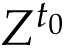and *θ* values.

### Data analysis

For all networks, we measured their connectance (the proportion of realised interactions from the pool of all possible interactions between the species of a network), number of components (the number of isolated groups of interacting species), and robustness to secondary extinctions. To assess robustness to secondary extinctions we progressively removed resource species at random and measured the proportion of consumer species that remain (Memmott et al., 2004). We quantified robustness as the area under the curve of the fraction of resources removed against the fraction of surviving consumers (Burgos et al., 2007). Thus, robustness can range between 0 (low robustness) and 1 (high robustness).

We compared the connectance, number of components, and robustness of networks built at the start of the simulation —when species traits were set to their random environmental optimum— and networks built after simulating coevolution for 200 generations —once traits had reached a steady state (see Figures S3-S5 for steady state values). The difference in these network properties can be attributed to the rewiring of networks due to coevolution.

We performed numerical simulations in Julia (version 1.4.2, Bezanson et al., 2017) and data analysis and visualisation in R (version 4.0.2, R Core Team, 2022).

## Results

Coevolution increases the robustness of mutualistic communities to secondary extinctions, but reduces it for antagonistic communities (Figure 2). In fact, coevolution reduces robustness in antagonism more than it increases it in mutualism. It does so by mediating the density of interactions in a community (Figure 3). While mutualistic coevolution boosts connectance, antagonistic coevolution decreases it. Hence, as coevolution builds up connectance, robustness increases as well. Yet, the exact relationship between connectance and robustness is shaped by community size and interaction type (Figure 4).

**Figure 2:**
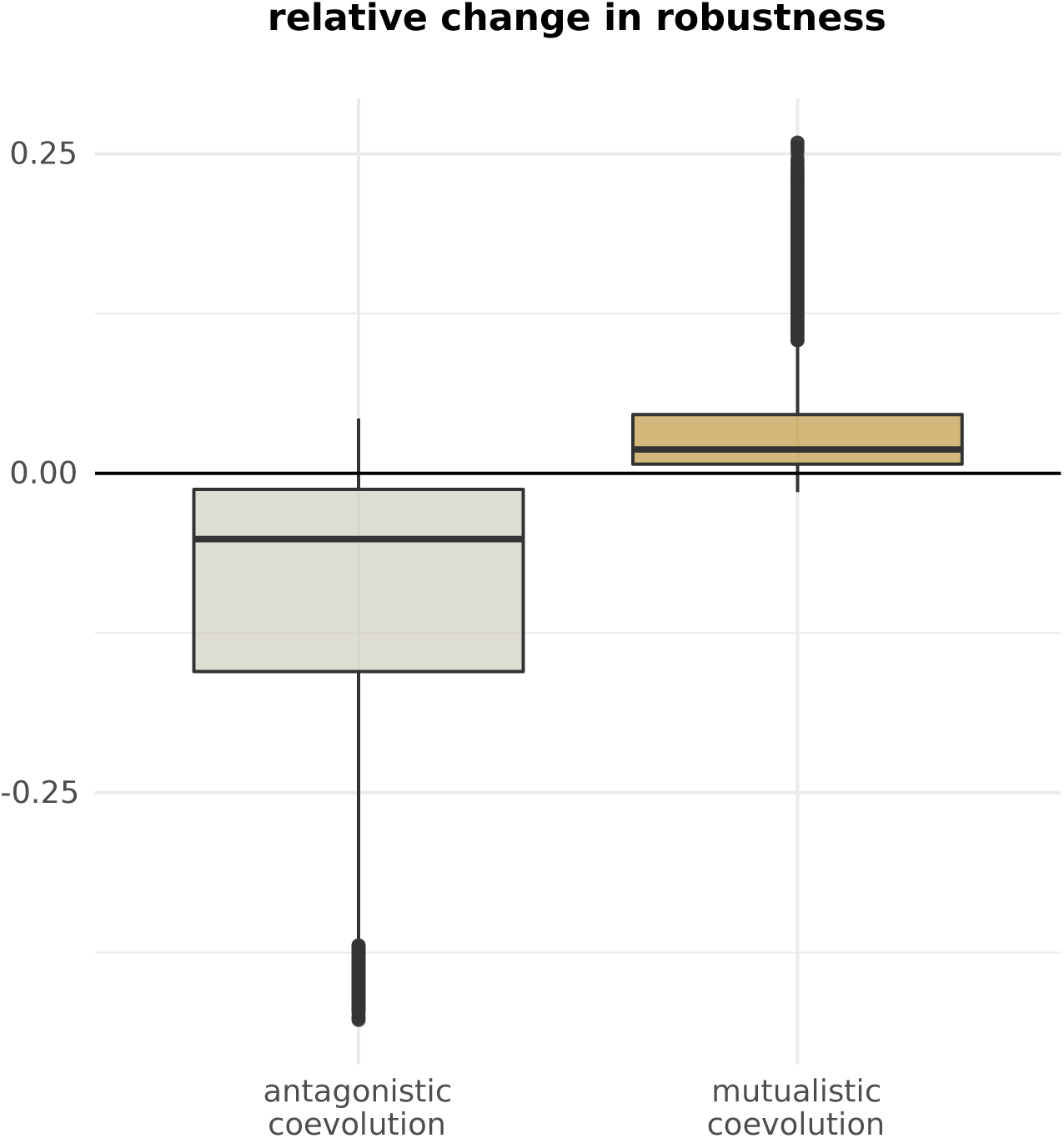
Coevolution alters the robustness to secondary extinctions. The boxplots summarise the relative change in network robustness after simulating coevolution across all communities and coevolutionary scenarios.

**Figure 3:**
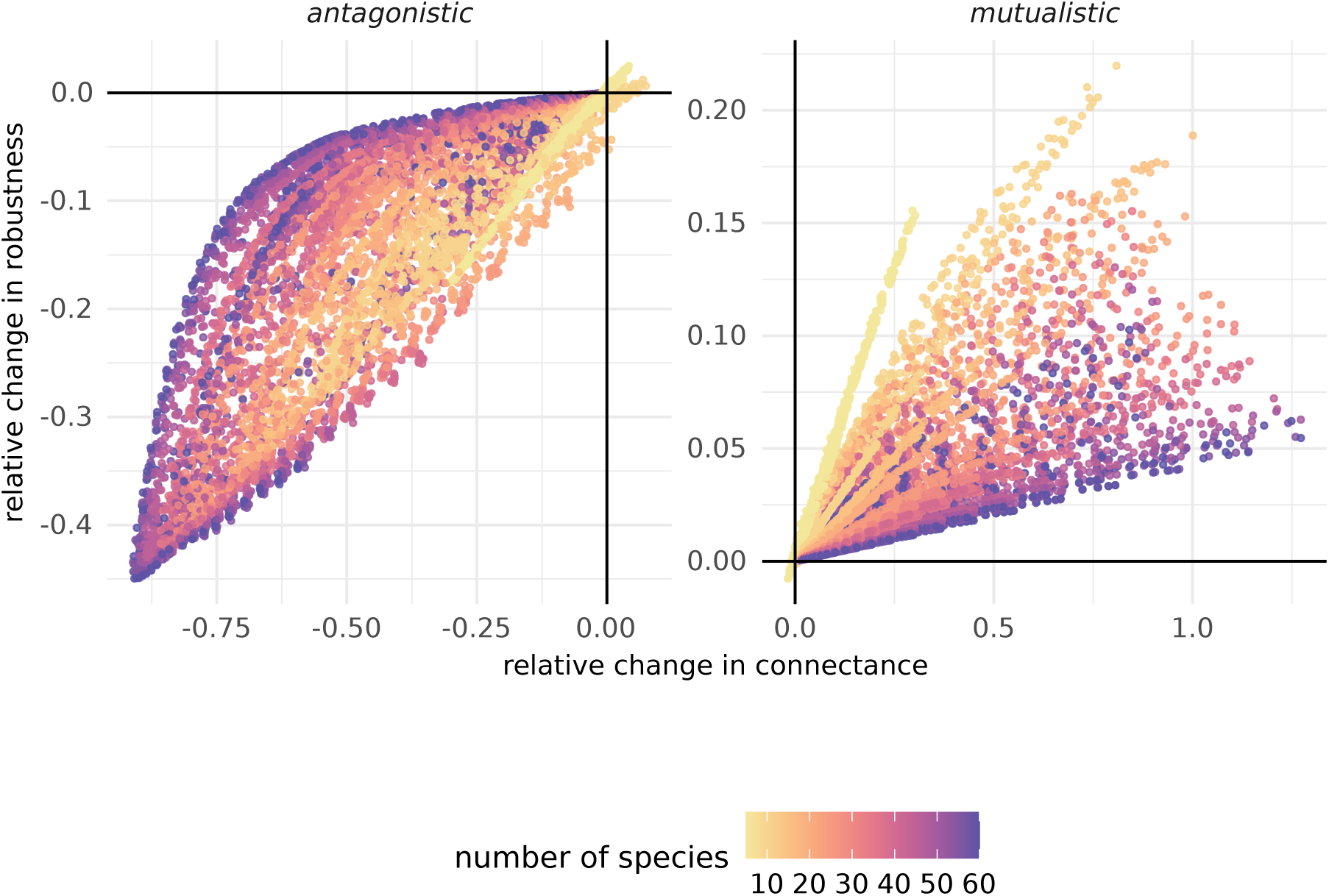
Coevolution shapes the robustness to secondary extinctions by altering network connectance. The figures show the changes in robustness and connectance of networks due to coevolution. Each point represents the average change in connectance and robustness for a given community under a particular coevolutionary scenario. The colour of points denotes the number of species in the community.

**Figure 4:**
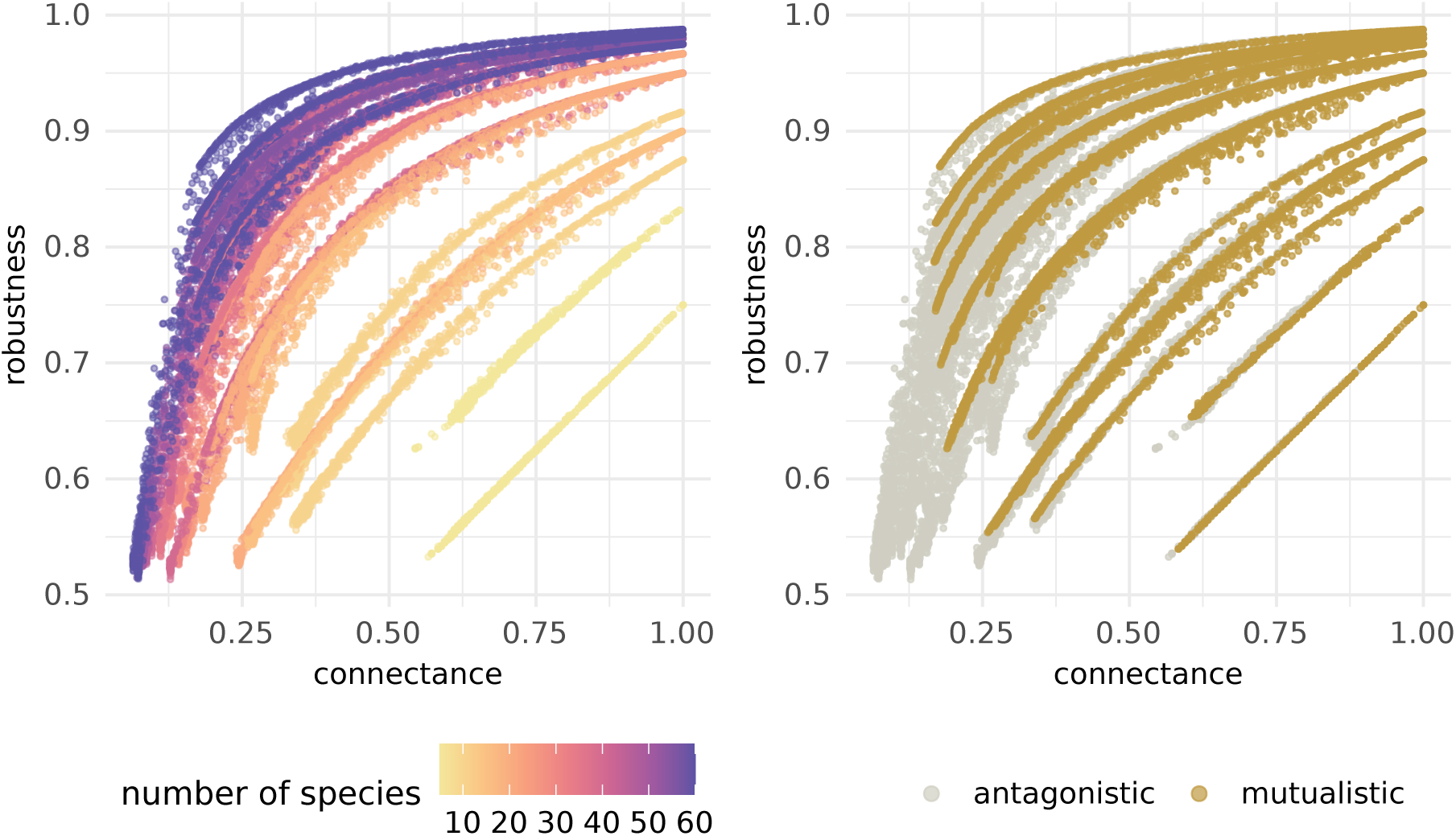
The community size and interaction type determine the relationship between connectance and robustness. Each point represents the average connectance and robustness for a given community under a particular coevolutionary scenario. In the left panel, the colour of points denotes the number of species in the community. In the right panel, the colour of points denotes the type of species interaction.

The relationship between robustness and connectance depends primarily on community size (Figure 4 left). The larger the community, the more non-linear the relationship. Nevertheless, the ratio of consumer to resource species in the community determines the exact shape of the relationship (Figure S11). Thus, each community has a particular connectance-robustness relationship (Figure S11). Yet, for a given community, different interaction types yield different regions of the robustness-connectance relationship (Figure 4 right). Mutualistic coevolution tends to build networks with high connectance and robustness (Figure 4 right, gold points). In contrast, the same communities, under antagonistic coevolution, tend to achieve lower connectance and robustness (Figure 4 right, silver points).

The consequence of a change in connectance on robustness differs between interaction types. The relationship between changes in connectance and robustness tends to be linear under mutualism and non-linear in antagonism (Figure 3). However, once more, the community size determines its exact shape. In mutualism, the slope of the linear relationship decreases with the size of the community (Figure 3 right). In antagonism, the relationship becomes increasingly non-linear with the size of the community (Figure 3 left). Thus, the type of species interactions and the size of a community influence the extent to which coevolution changes robustness.

The conditions under which coevolution operates modulate the size of changes to robustness and connectance (Figure 5). The largest changes occur when both coevolutionary selection and functional mechanism mediating coevolution are strong (Figure 5, high *m* and *α* values). This is true for both mutualism and antagonism. Yet, the direction of changes differs between the two.

**Figure 5:**
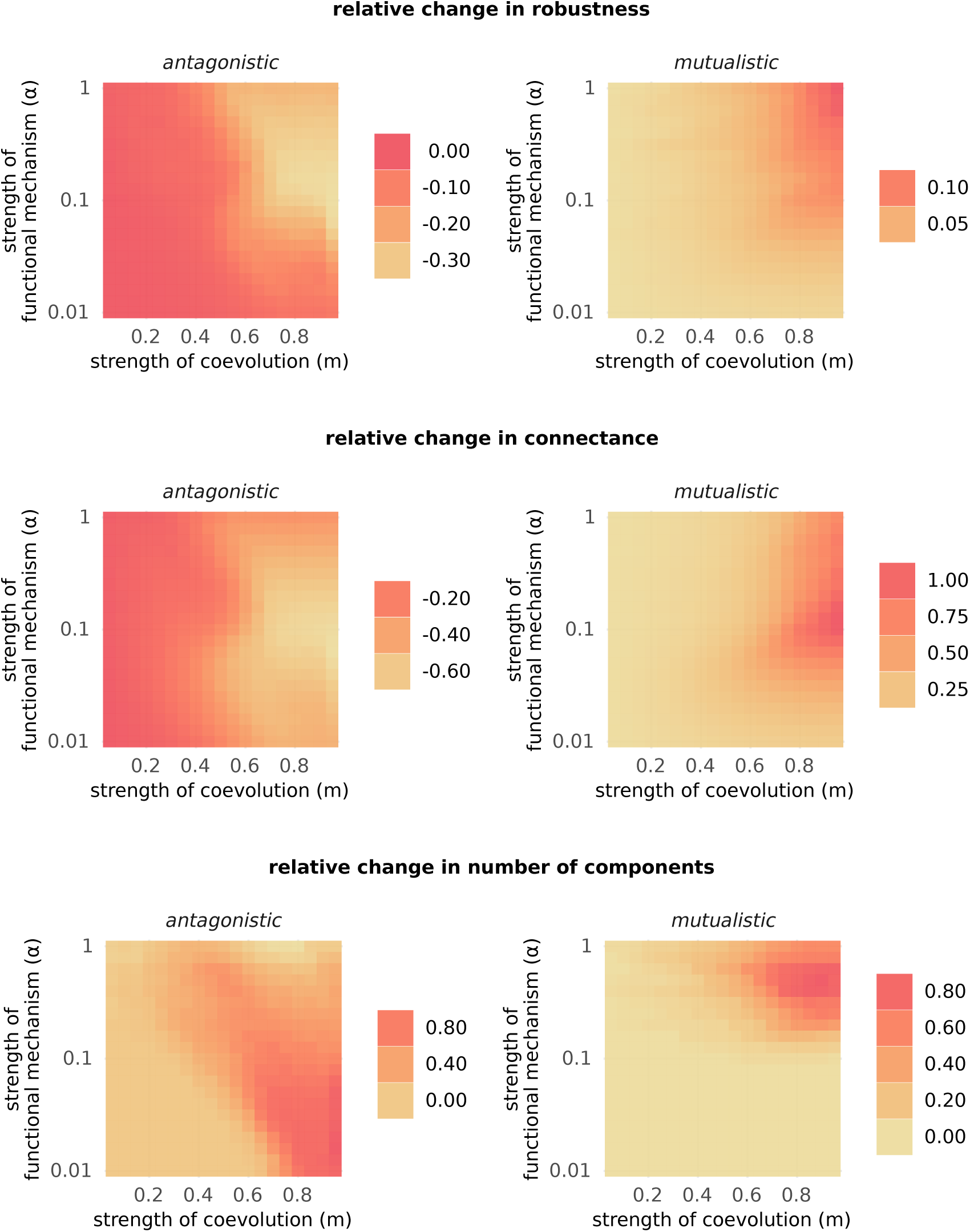
The strength of coevolution and functional mechanism controls the magnitude and direction of changes to robustness, connectance, and number of components. The grids summarise the relative change in robustness, connectance, or number of components due to coevolution, averaged across all communities for each coevolutionary scenario.

Coevolution also influences the number of components in a network (Figures 5 and S5) which, in turn, affects its connectance and robustness (Figure S6). Mutualistic coevolution tends to maintain networks with a single component (Figures 5 and S5). However, antagonistic coevolution tends to fragment networks into many components (Figures 5 and S5). In mutualism, fragmentation into components happens under strong coevolutionary selection and functional mechanism (Figure 5). Still, even when fragmented, mutualistic networks increase their connectance, and robustness (Figures 5 and S6). In antagonism, the relationship between the changes in the number of components, connectance and robustness depends on the coevolutionary scenario. For example, under strong coevolutionary selection, a weak or strong functional mechanism yields similar changes to connectance (Figure 5, high *m* values). But networks fragment more when the functional mechanism is weak (Figure 5, high *m* and low *α* values). This leads to smaller reductions in robustness than when the functional mechanism is strong.

## Discussion

We find that coevolution has the potential to affect the robustness of communities to secondary extinctions. These changes in robustness are due to changes in the structure of interaction networks. Yet, the impact of coevolution on robustness is not the same across all communities. First, the type of species interactions determines the direction of change in robustness. Mutualistic coevolution increases robustness while antagonistic coevolution reduces it. Second, the intensity of coevolution modulates the magnitude of changes in robustness. The largest changes in robustness occur under strong coevolution. Third, the size of communities impacts the size of changes to robustness. Small mutualistic communities experience the largest gains in robustness, while large antagonistic communities have the biggest losses.

Coevolution can lead to changes in the traits of interacting species (Thompson, 2005). These changes in traits can in turn affect the establishment and maintenance of interactions (Thompson, 2005; Santamaŕıa and Rodŕıguez-Gironés, 2007; Vázquez et al., 2009; McPeek, 2017; Guimarães, 2020). In this sense, we find that coevolution can affect the density of interactions in communities. Yet, it does so differently in mutualistic and antagonistic assemblages (Figure 2). This is due to the fact that mutualism increases trait complementarity between partners while antagonism reduces it. So, coevolution increases the density of interactions in mutualism but decreases it in antagonism (Figure 2). Crucially, we show that these changes in connectance also affect the robustness of communities. In agreement with previous work, we find a positive association between connectance and robustness (Dunne et al., 2002; Eklö f and Ebenman, 2006).

The nature of species interactions may influence the intensity of coevolutionary selection (Thompson, 2005). For instance, selective pressures may be stronger in obligate rather than facultative interactions (Thompson, 1994). Moreover, some interactions may require partners to have similar traits in order to interact. For example, flower size can constrain the morphology of potential pollinators (Stang et al., 2006). We tested how these different coevolutionary scenarios affected the structure and robustness of communities. In agreement with Nuismer et al. (2013), strong coevolution leads to a high density of mutualistic interactions (Figure 5). In contrast, strong coevolution leads to a low density of antagonistic interactions (Figure 5). Consequently, the largest changes to robustness occur under strong coevolution (Figure 5). Therefore, the effect of coevolution on robustness may depend on the nature of species interactions.

A network can transition from a connected to a disconnected state depending on the distribution of interactions per species (Newman et al., 2001; Guimarães, 2020). Factors such as the intimacy of interactions or the heterogeneity of traits that mediate interactions may influence whether a network is connected or not (Guimarães, 2020). We show that coevolution can also shift communities from one state to the other (Figures 5 and S5). Disconnected networks are more common when coevolutionary selection is strong and when communities are antagonistic (Figure 5). In antagonism, depending on the coevolutionary scenario, fragmented components may either increase or reduce robustness. Yet, even when fragmented, mutualistic networks retain a high density of interactions and are more robust than antagonistic communities (Figure S6). In summary, coevolution can influence whether networks are connected or disconnected. This, in turn, impacts the spread of co-extinctions.

The effect of coevolution on robustness may depend on the size of the community. As noted above, we observe a positive association between connectance and robustness. Yet, we find the shape of this relationship depends on species richness (Figure 4). Under mutualistic coevolution, small communities undergo the largest gains in robustness (Figure 3 right). Under antagonistic coevolution, large communities experience non-linear losses in robustness (Figure 3 left). These results strengthen the argument that the size of communities affects the propensity of co-extinctions (Vanbergen et al., 2017). Moreover, they suggest that potential beneficial or detrimental effects of coevolution on robustness could be amplified by the size of the community.

As with any modelling approach, our findings are contingent on our assumptions. In particular, we identify three that could impact the outcome of our results. First, we assumed that species interactions lead to trait matching in mutualism and trait mismatch in antagonism. Although this is a reasonable assumption, evidence suggests that traits can evolve in other ways (e.g., Alexandersson and Johnson, 2002; Nuismer and Thompson, 2006). Different functional mechanisms, such as trait barriers, will likely lead to different dynamics (Nuismer and Thompson, 2006; Andreazzi et al., 2020). An extension of our work could test the generality of our findings under different mechanisms of trait evolution. Second, to build interaction networks, we converted the probabilities of interactions into “success” or “failure” events by drawing random numbers. Thus, each network realisation depends, to an extent, on the particular random variable being drawn. To minimise the effect of chance, one could treat the matrix of probabilities of interactions as a surrogate for interaction strength. This could be particularly relevant as interaction strength has also been shown to affect robustness (Kaiser-Bunbury et al., 2010; Gaiarsa and Guimarães, 2019). Third, we assumed that co-extinctions occurred only after coevolution had stopped operating once species traits had reached an equilibrium. Moreover, we did not allow communities to further coevolve throughout the co-extinction simulations. Yet, ecological and evolutionary processes can operate at similar timescales and influence each other (Carroll et al., 2007). Future work could test how coevolution affects the robustness of communities whilst co-extinctions are ongoing.

Extinction events have become increasingly common (Ceballos et al., 2015). This loss of biodiversity could be further amplified by co-extinction cascades (Cowie et al., 2022). As such, species interactions play a crucial role in the maintenance of biodiversity. Networks of interactions could either exacerbate or mitigate the loss of biodiversity. To inform conservation efforts, we must have an understanding of the factors that affect the robustness of communities (Bascompte and Scheffer, 2023). We argue that coevolution may be one such factor. Thus, fostering or limiting the coevolution of populations may serve as a mechanism for increasing robustness. Naturally, these insights must first be empirically tested. This undoubtedly is a major challenge. Yet, the current extinction crisis calls for a deep understanding of the processes that mitigate co-extinctions.

## Author contributions

FP, KAG, and JB designed research. FP performed research and together with KAG analysed data. FP wrote a first draft of the manuscript, and all authors contributed substantially to the final draft.

## Acknowledgements

We thank the members of Bascompte Lab and Cecilia Andreazzi for discussions. Funding was provided by the University of Zurich Research Priority Program Global Change and Biodiversity (URPP GCB), University of Zurich Postdoc Grant (grant number FK-22-114 to KAG), and the SNSF (grant number 310030 197201 to JB).

## Code and data availability

All code and data is available at https://github.com/fp3draza/coevolution_robustness and https://github.com/fp3draza/coevolution_robustness/releases respectively.

---

**Figure S1:**
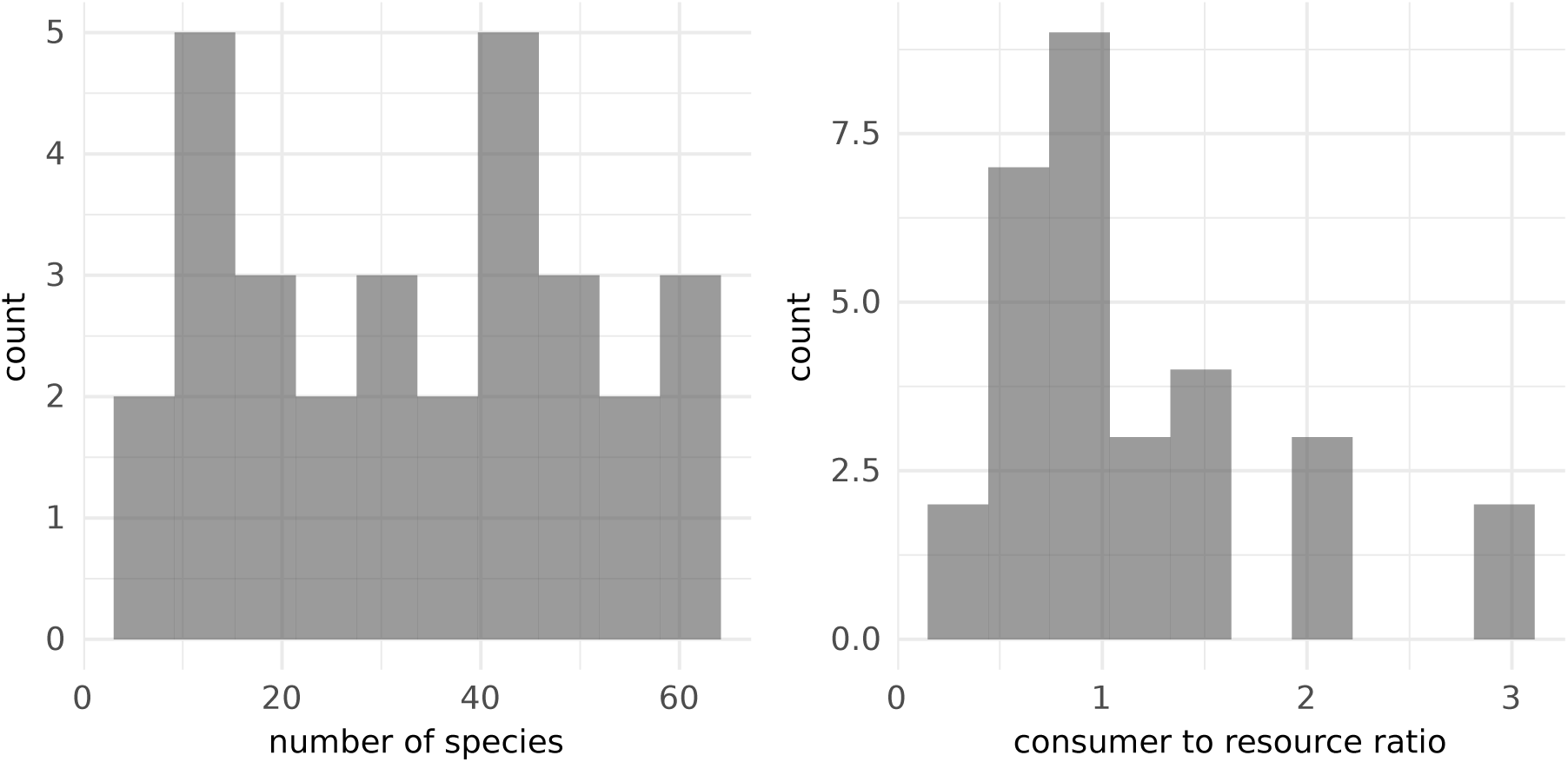
Size and ratio of consumers to resources of communities used to parametrise simulations.

**Figure S2:**
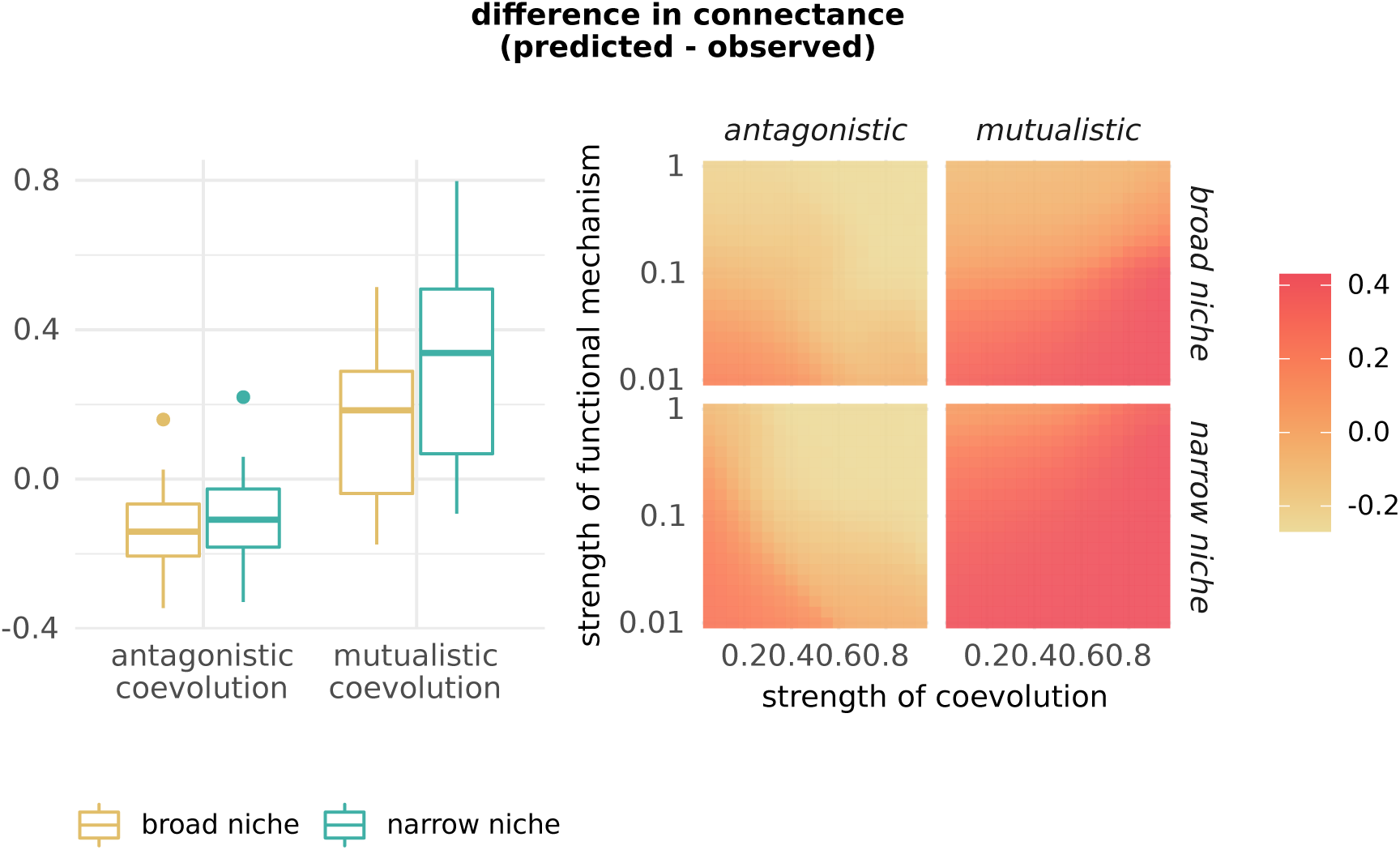
Validation of the modelling framework. The left panel shows the difference between predicted and observed connectance values of a set of empirical antagonistic and mutualistic networks. The right panel summarises the average difference between predicted and observed connectance values, in different coevolutionary scenarios.

**Figure S3:**
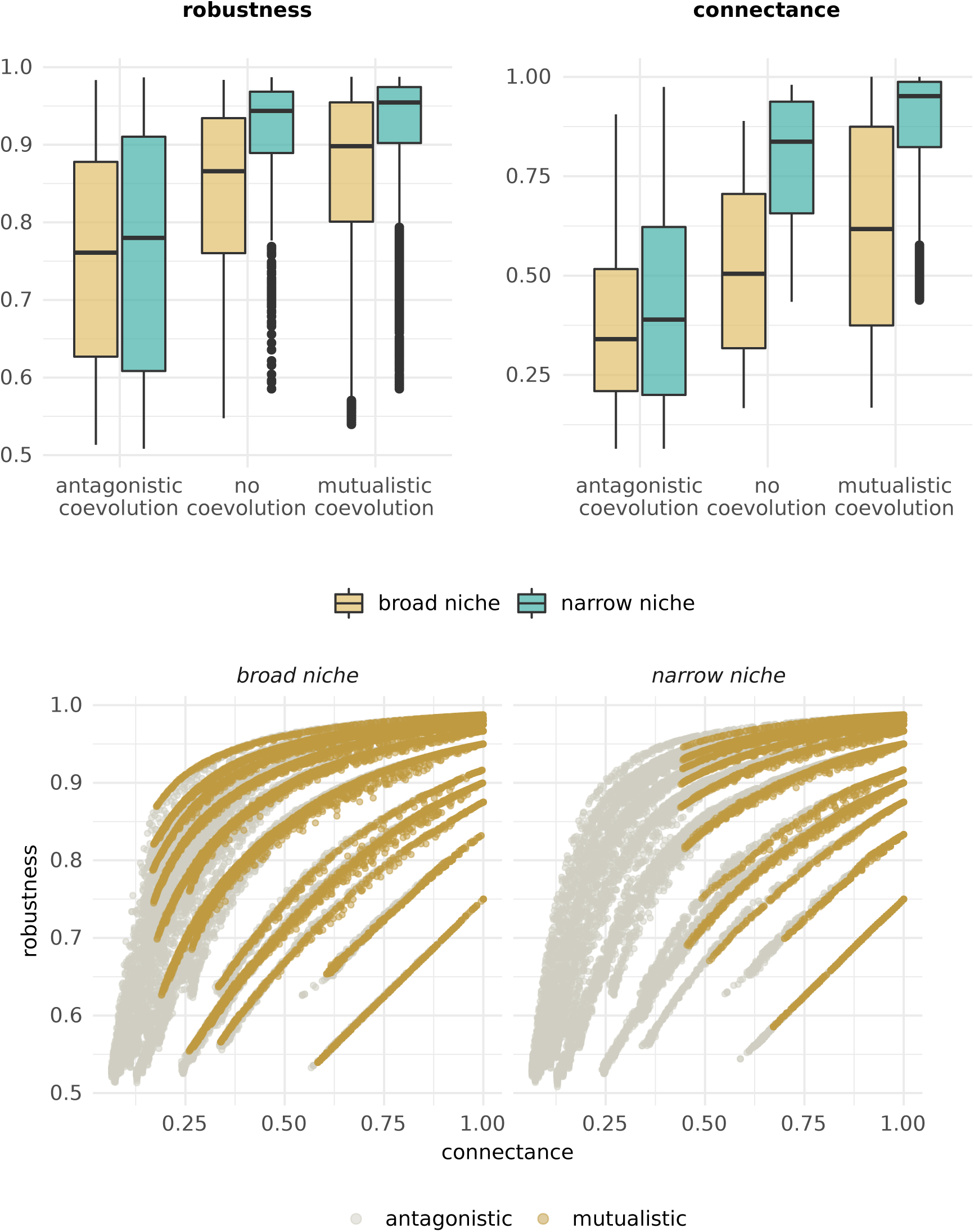
Coevolution shapes the robustness to secondary extinctions by altering network connectance. The top row compares the network robustness and connectance of communities under: antagonistic coevolution, no coevolution (i.e. when species traits are fixed to their environmental optimum), and mutualistic coevolution. The bottom row shows the relationship between network connectance and robustness after coevolution. Each point represents the average connectance and robustness of a given community under a particular coevolutionary scenario.

**Figure S4:**
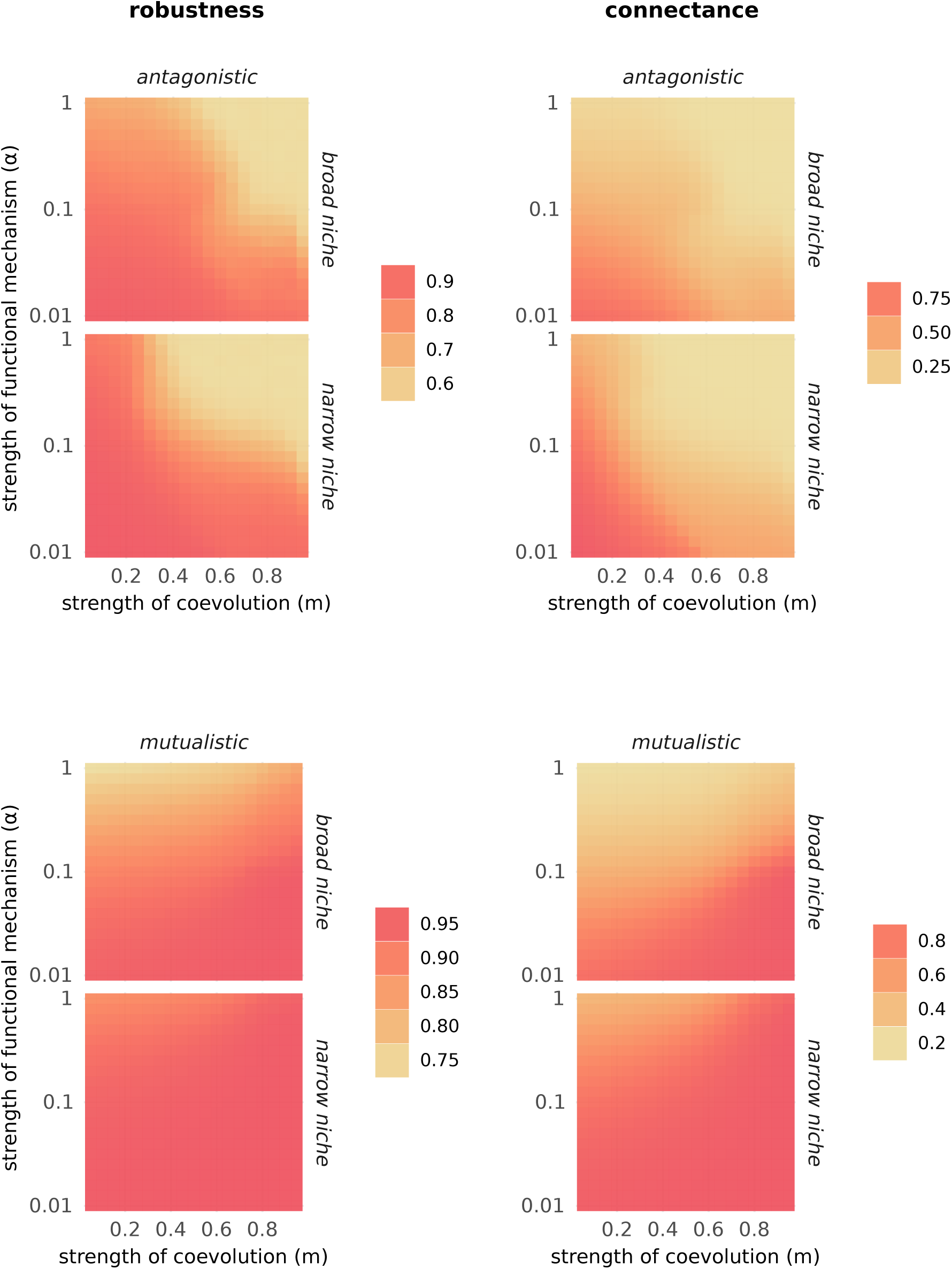
The strength of coevolution controls robustness and connectance. The grids summarise the average robustness or connectance after coevolution, in different coevolutionary scenarios.

**Figure S5:**
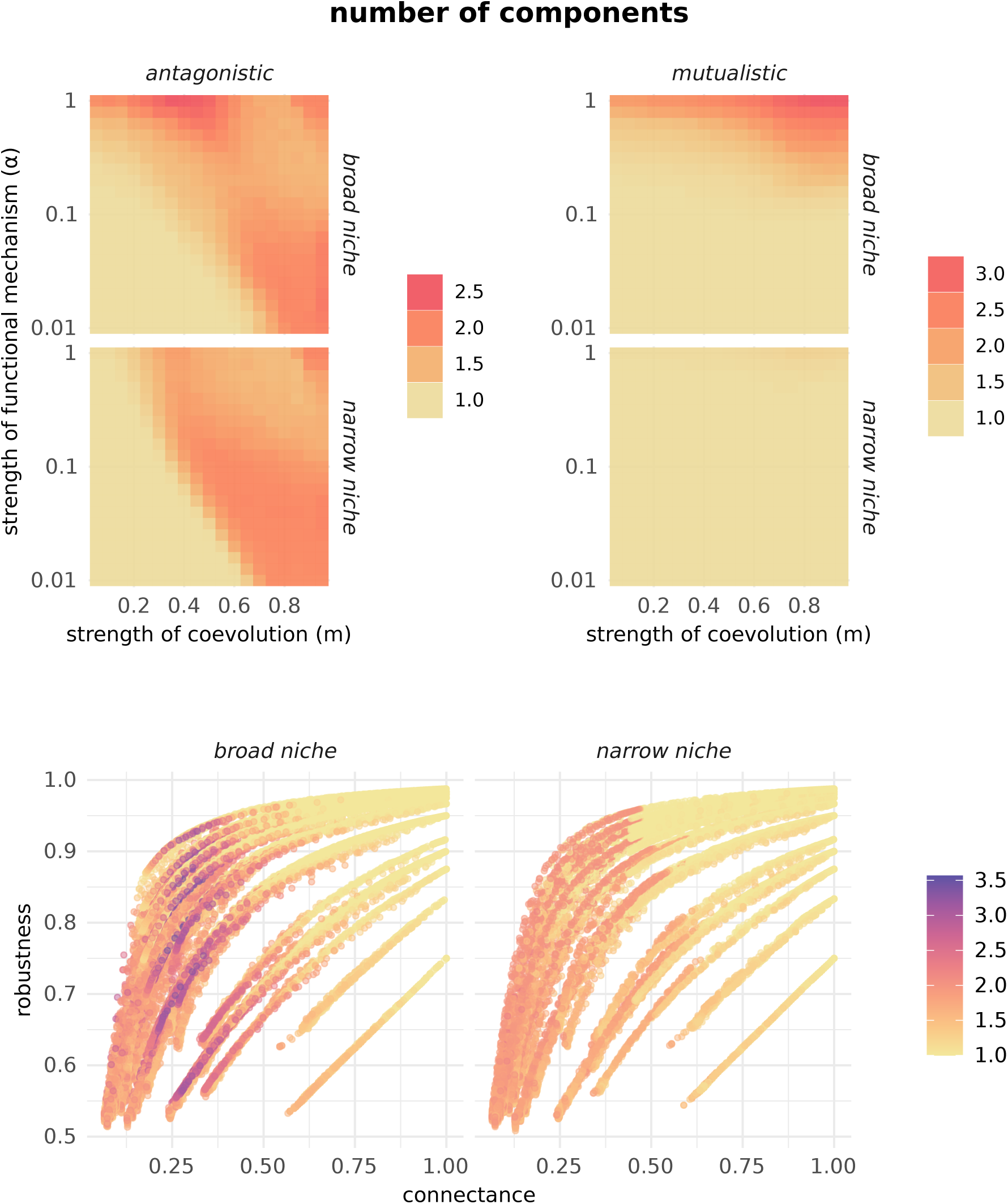
The number of components in a community influences its robustness and connectance. The grids summarise the average number of components in communities after coevolution, in different coevolutionary scenarios. The bottom row shows the relationship between network connectance and robustness after coevolution. Each point represents the average connectance and robustness for a given community under a particular coevolutionary scenario. The colour of points denotes the average number of components.

**Figure S6:**
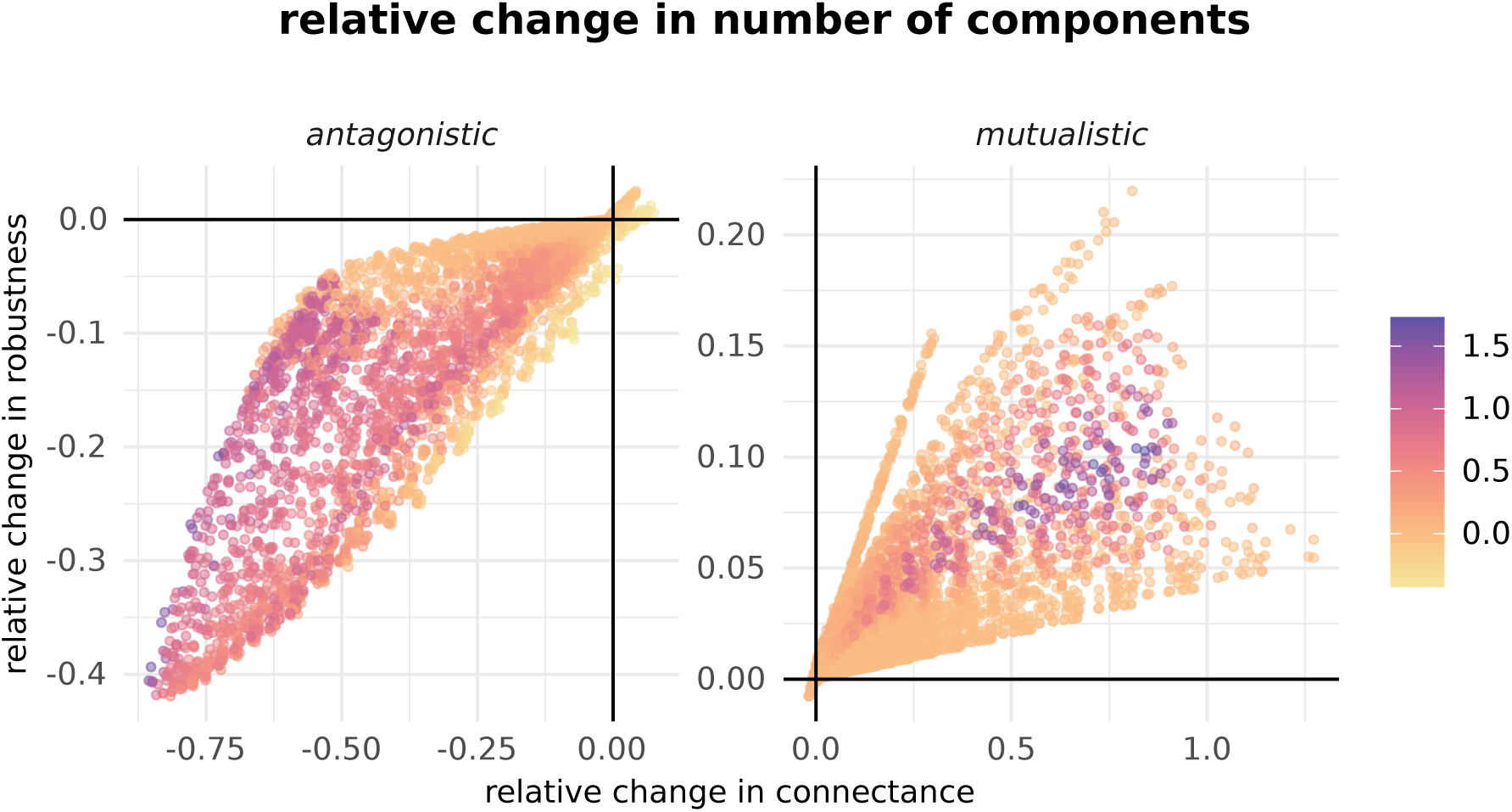
The number of components in a community influences its connectance and robustness. The figure shows the relationship between changes in network connectance and robustness after coevolution. Each point represents the average change in connectance and robustness for a given community under a particular coevolutionary scenario. The colour of points denotes the average relative change in number of components.

**Figure S7:**
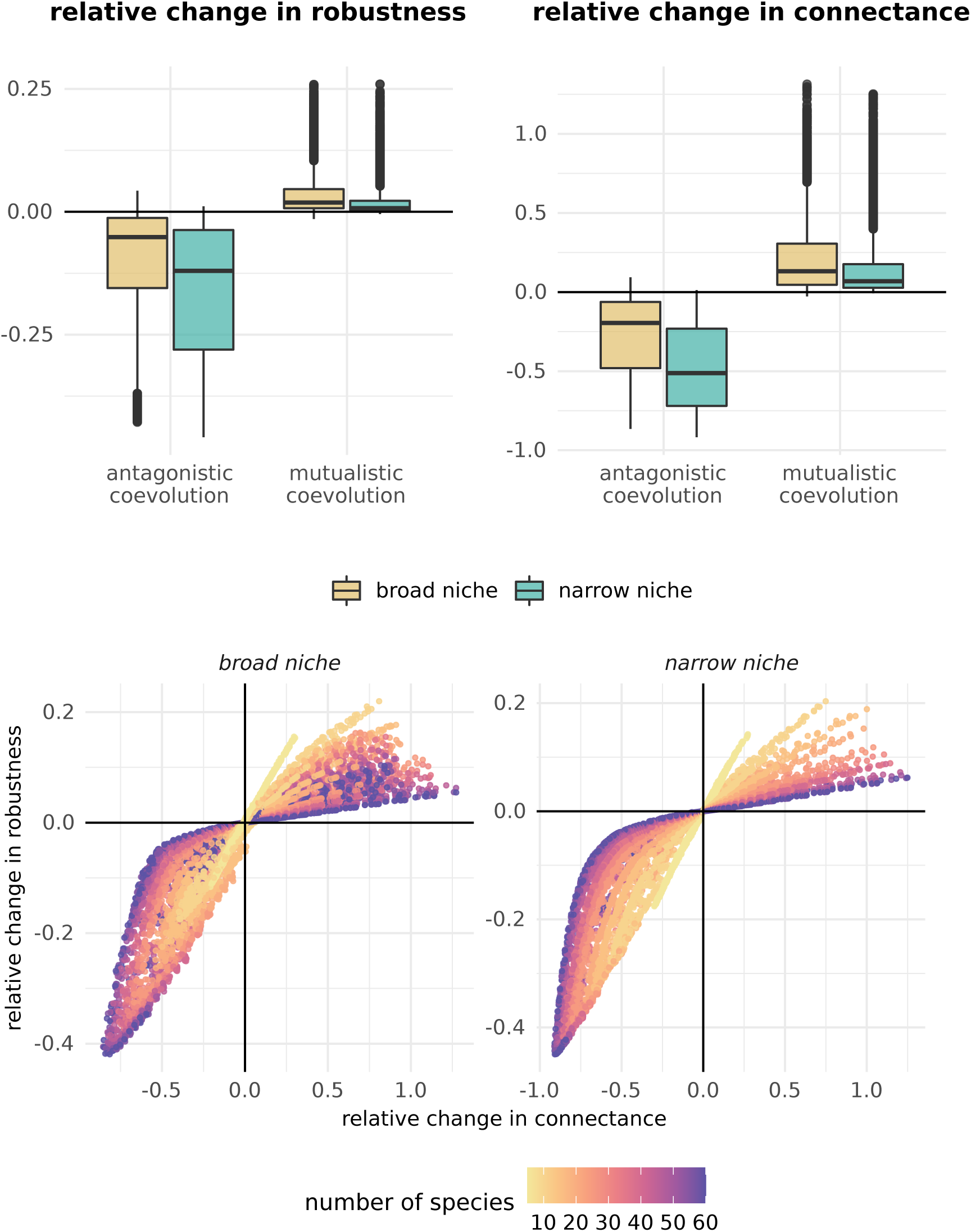
Coevolution shapes the robustness to secondary extinctions by altering network connectance. The top row summarises the relative change in network robustness and connectance after simulating coevolution across all communities. The bottom row shows the relationship between changes in network connectance and robustness after coevolution. Each point represents the average change in connectance and robustness for a given community under a particular coevolutionary scenario.

**Figure S8:**
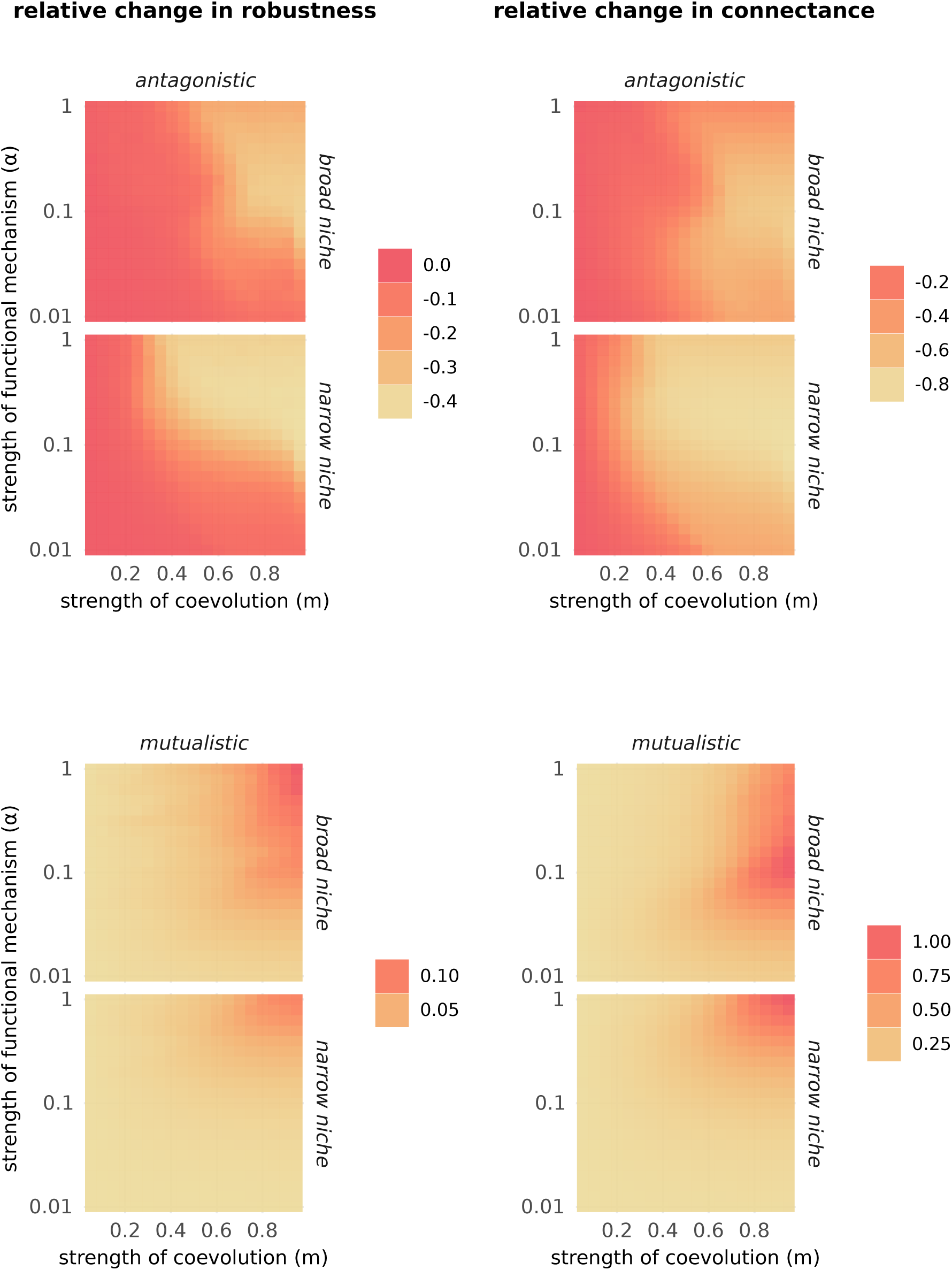
The strength of coevolution and functional mechanism controls the magnitude and direction of changes to robustness and connectance. The grids summarise the average relative change in robustness or connectance due to coevolution, across all communties for each coevolutionary scenario.

**Figure S9:**
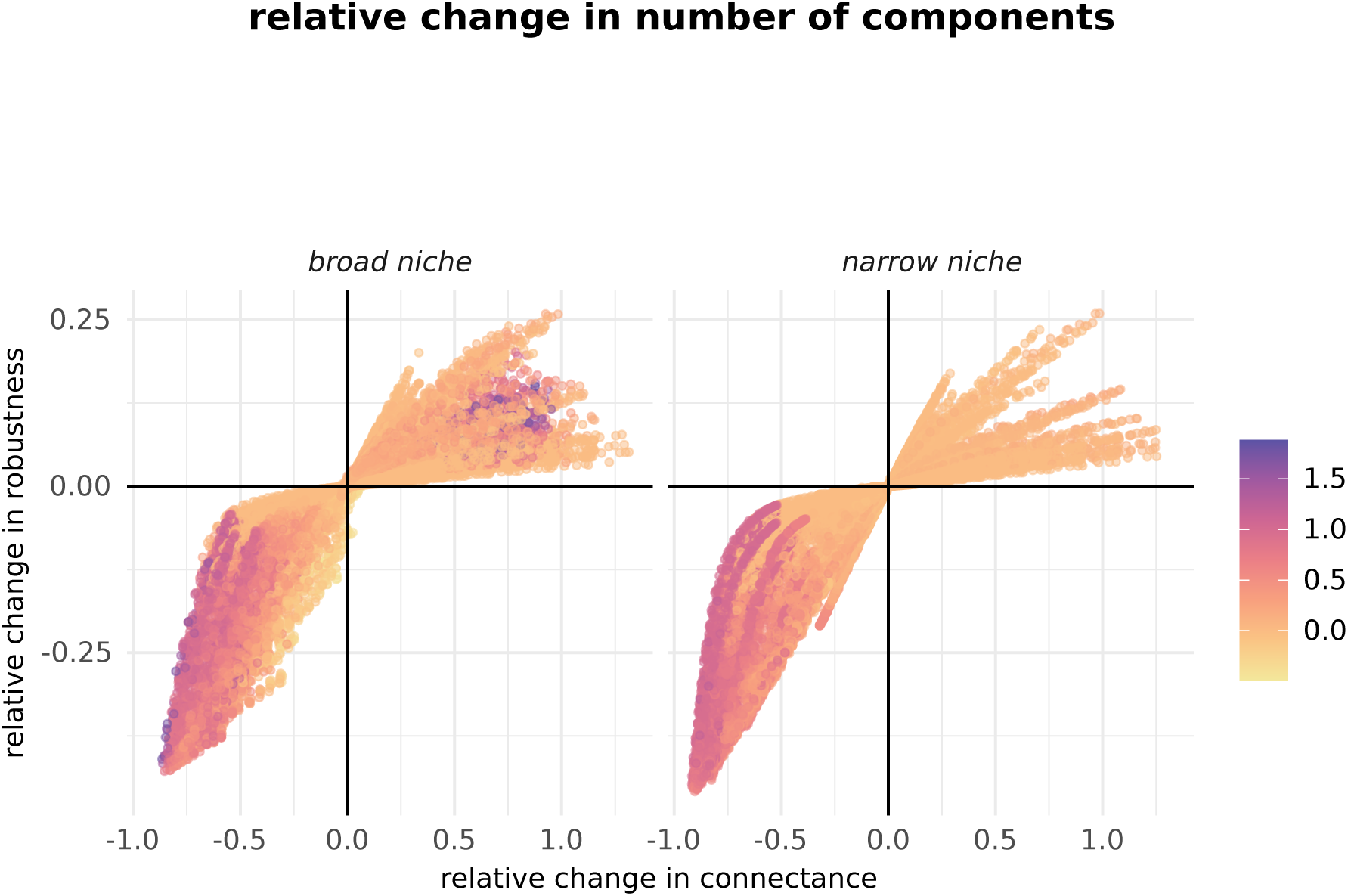
The number of components in a community influences its connectance and robustness. The grids summarise the average relative change in the number of components in communities due to coevolution, across all communities. The bottom row shows the relationship between changes in network connectance and robustness after coevolution. Each point represents the average change in connectance and robustness for a given community under a particular coevolutionary scenario. The colour of points denotes the average relative change in number of components.

**Figure S10:**
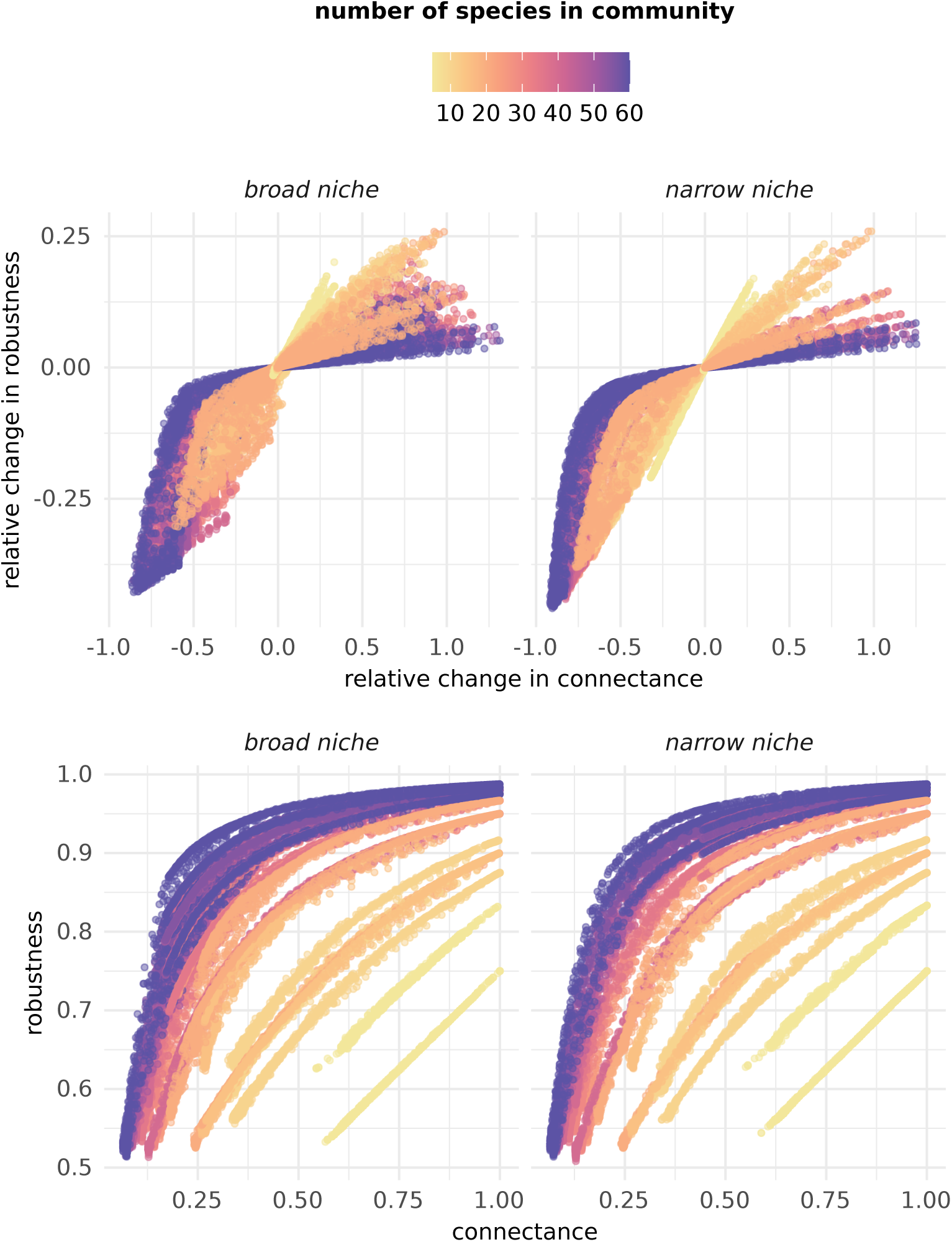
The number of species in a community modulates how coevolution shapes network robustness and connectance. The top row shows the relationship between the relative change in network connectance and robustness due to coevolution. The bottom row shows the relationship between network connectance and robustness after coevolution. Each point represents the average connectance and robustness for a given community under a particular coevolutionary scenario. The colour of points denotes the number of species in each community.

**Figure S11:**
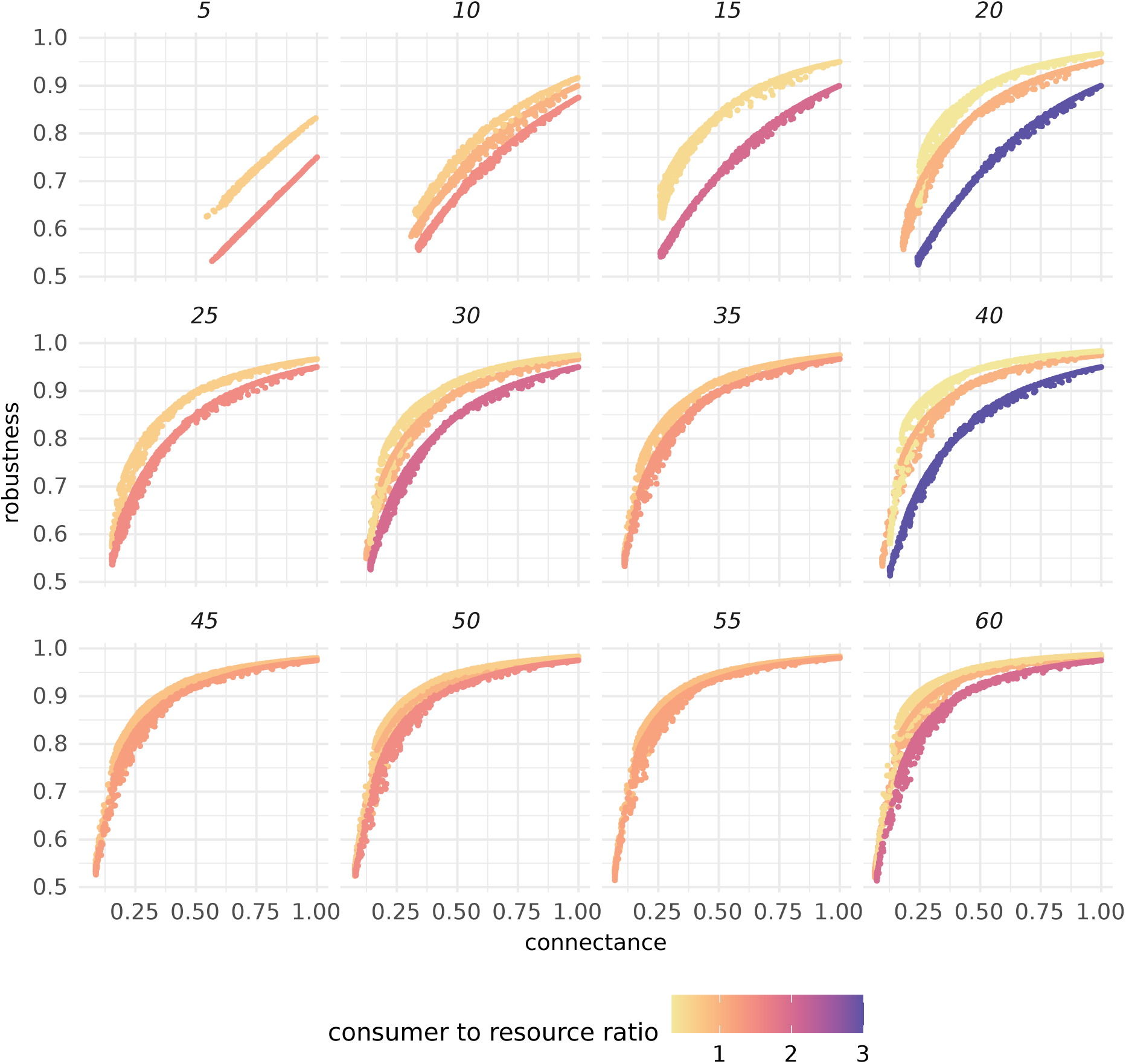
The effect of the ratio of consumer to resource species in shaping the relationship between connectance and robustness. The number above each panel denotes the number of species in the community. Each point represents the average connectance and robustness for a given community under a particular coevolutionary scenario. The colour of points denotes the ratio of consumer to resource species in the community.

**Figure S12:**
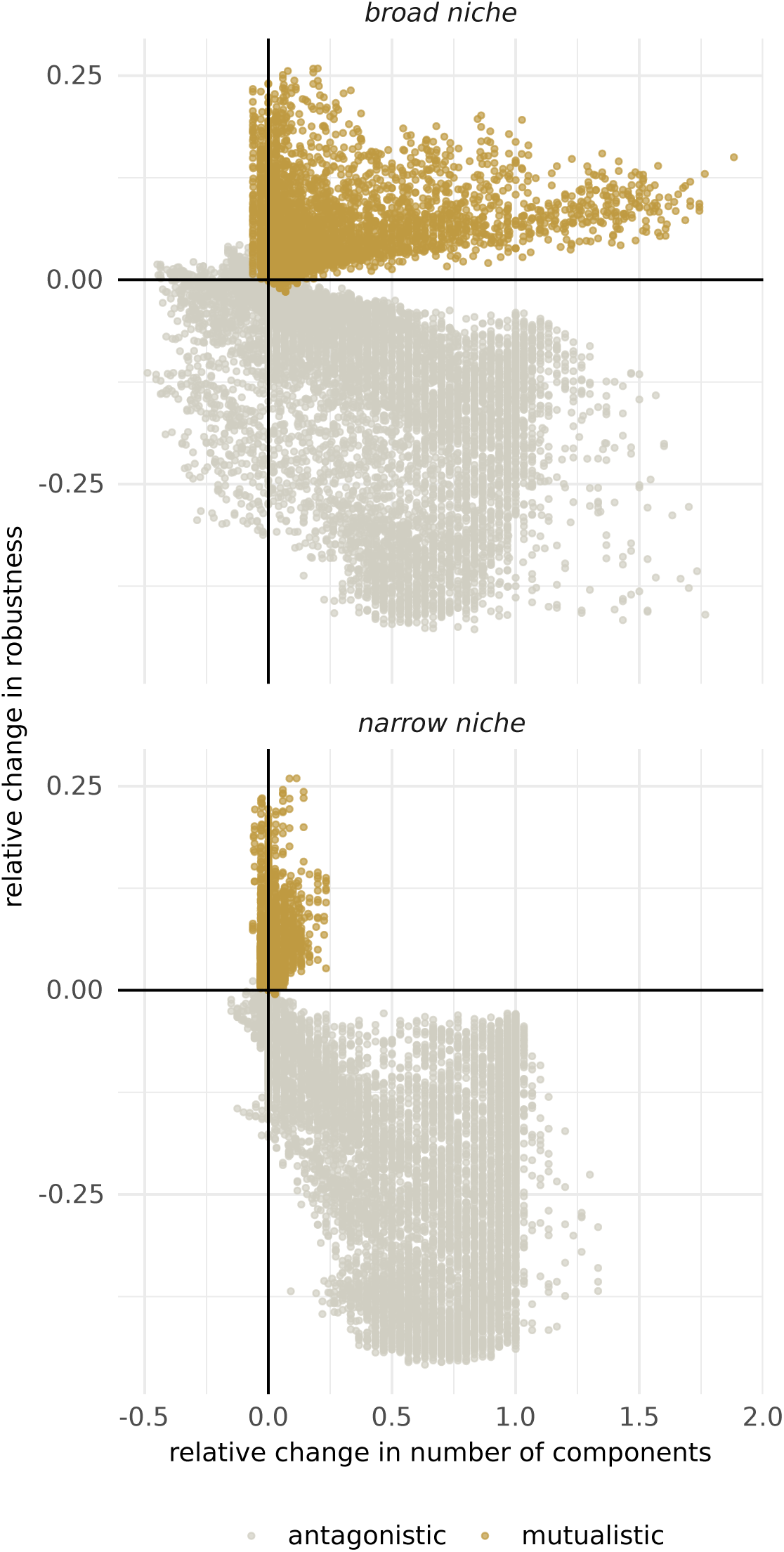
The type of species interactions modulates the extent to which coevolution fragments networks into components and their robustness. Each point represents the average change in the number of components and change in robustness for a given community under a particular coevolutionary scenario. The colour of points denotes the type of species interaction.

